# Production and cross-feeding of nitrite within *Prochlorococcus* populations

**DOI:** 10.1101/2021.12.31.474641

**Authors:** Paul M. Berube, Tyler O’Keefe, Anna Rasmussen, Sallie W. Chisholm

## Abstract

*Prochlorococcus* is an abundant photosynthetic bacterium in the oligotrophic open ocean where nitrogen (N) often limits the growth of phytoplankton. *Prochlorococcus* has evolved into multiple phylogenetic clades of high-light (HL) adapted and low-light (LL) adapted cells. Within these clades, cells encode a variety of N assimilation traits that are differentially distributed among members of the population. Among these traits, nitrate (NO_3_^−^) assimilation is generally restricted to a few clades of high-light adapted cells (the HLI, HLII, and HLVI clades) and a single clade of low-light adapted cells (the LLI clade). Most, if not all, cells belonging to the LLI clade have the ability to assimilate nitrite (NO2^−^), with a subset of this clade capable of assimilating both NO_3_^−^ and NO_2_^−^. Cells belonging to the LLI clade are maximally abundant at the top of the nitracline and near the primary NO_2_^−^ maximum layer. In some ecosystems, this peak in NO_2_^−^ concentration may be a consequence of incomplete assimilatory NO_3_^−^ reduction by phytoplankton. This phenomenon is characterized by a bottleneck in the downstream half of the NO_3_^−^ assimilation pathway and the concomitant accumulation and release of NO_2_^−^ by phytoplankton cells. Given the association between LLI *Prochlorococcus* and the primary NO_2_^−^ maximum layer, we hypothesized that some *Prochlorococcus* exhibit incomplete assimilatory NO_3_^−^ reduction. To assess this, we monitored NO_2_^−^ accumulation in batch culture for 3 *Prochlorococcus* strains (MIT0915, MIT0917, and SB) and 2 *Synechococcus* strains (WH8102 and WH7803) when grown on NO_3_^−^ as the sole N source. Only MIT0917 and SB accumulated external NO_2_^−^ during growth on NO_3_^−^. Approximately 20-30% of the NO_3_-transported into the cell by MIT0917 was released as NO_2_^−^, with the balance assimilated into biomass. We further observed that co-cultures using NO_3_- as the sole N source could be established for MIT0917 and a *Prochlorococcus* strain that can assimilate NO_2_^−^ but not NO_3_^−^. In these co-cultures, the NO_2_^−^ released by MIT0917 was efficiently consumed by its partner strain during balanced exponential growth. Our findings highlight the potential for emergent metabolic partnerships within *Prochlorococcus* populations that are mediated by the production and consumption of the N cycle intermediate, NO_2_^−^.

**SIGNIFICANCE:** Earth’s biogeochemical cycles are substantially driven by microorganisms and their interactions. Given that N often limits marine photosynthesis, we investigated the potential for N cross-feeding within populations of *Prochlorococcus*, the numerically dominant photosynthetic cell in the subtropical open ocean. During growth on NO_3_^−^, some *Prochlorococcus* cells release up to 30% of their N uptake as extracellular NO_2_^−^. In the wild, *Prochlorococcus* populations are composed of multiple functional types, including those that cannot use NO_3_^−^ but can still assimilate NO_2_^−^. We show that metabolic dependencies arise when *Prochlorococcus* strains with complementary NO_2_^−^ production and consumption phenotypes are grown together on NO_3_^−^. These findings demonstrate the potential for emergent metabolic partnerships, possibly modulating ocean nutrient gradients, that are mediated by cross-feeding of N cycle intermediates.

## INTRODUCTION

*Prochlorococcus* and its close relative, *Synechococcus*, are globally abundant cyanobacteria that are jointly responsible for approximately 25% of marine net primary production – roughly 12 gigatons of fixed carbon each year [1]. The share of marine primary production attributed to *Prochlorococcus* is predicted to increase over the course of this century as a consequence of a warmer and more stratified ocean [1, 2]. Phytoplankton growth, and thus primary production, is limited by N across much of the surface ocean [3]. Given the numerical dominance of *Prochlorococcus*, examining its genotypic and phenotypic diversity in the context of the N cycle can inform our understanding of *Prochlorococcus’* role in marine primary production.

*Prochlorococcus* has multiple N assimilation traits, most of which are distributed across cells in *Prochlorococcus* populations. All *Prochlorococcus* appear capable of assimilating ammonium (NH_4_^+^), generally the preferred N source for cyanobacteria since it is readily incorporated into the amino acid pool [4]. Some *Prochlorococcus* also contain flexible genes – found in some, but not all genomes – that enable the assimilation of organic N sources such as urea, cyanate, and amino acids [5, 6, 7] as well as the inorganic N sources, NO_3_^−^ and NO_2_^−^ [8, 9, 10, 11]. NO_3_^−^ assimilation by *Prochlorococcus* was identified decades after the first strain of *Prochlorococcus* was brought into culture [8, 9, 12] – prior to this discovery, it was thought that *Prochlorococcus* lacked this functional trait because isolated cultures could not use NO3^−^ as a N source and NO_3_^−^ assimilation genes were absent among the first sequenced genomes [6, 13]. It is now evident that there is extensive variability with respect to the N assimilation traits harbored by *Prochlorococcus*. These biological features have the potential to impact N cycling across the vast subtropical ocean gyres – the consequences of which are not well constrained.

The evolutionary history of NO_3_^−^ assimilation in *Prochlorococcus* has deepened our understanding of the selective pressures operating on this organism. It appears likely that the NO_3_^−^ assimilation gene cluster has been present in *Prochlorococcus* since their divergence from *Synechococcus*, but only retained in a subset of more recently emerged clades [11]. While loss of the NO_3_^−^ assimilation trait has essentially run to completion in most low-light adapted clades, the trait has been retained in a monophyletic group of high-light adapted clades as well as the LLI clade [11]. High-light adapted cells with NO_3_^−^ assimilation genes appear to be selected for in the surface waters of N-limited systems [10]. In contrast, selection for LLI *Prochlorococcus* cells with the capacity to assimilate NO_3_^−^ is not well resolved. This group of *Prochlorococcus* is widely distributed and abundant, often exceeding the combined depth-integrated abundance of other low-light adapted *Prochlorococcus* [14, 15]. LLI cells tolerate extreme fluctuations in light intensity allowing them to survive in surface waters where most low-light adapted *Prochlorococcus* are essentially absent [15, 16]. Cells belonging to the LLI clade typically have highest cell abundances in the subsurface chlorophyll maximum layer where nutrient concentrations are higher than the mixed layer [17, 18]. Intriguingly, we observed a spatial relationship between LLI cells with the NO_3_^−^ assimilation trait and a peak in NO_2_^−^ concentration in the water column [10].

In stratified marine systems, elevated concentrations of NO_2_^−^ in the mid-euphotic zone are ubiquitous. This feature, the primary NO_2_^−^ maximum layer, is thought to arise from distinct biological processes – either decoupled nitrification or incomplete NO_3_^−^ reduction by phytoplankton [19]. The respective activities of ammonia-oxidizing and nitrite-oxidizing microorganisms – performing the sequential reactions of nitrification – could drive the accumulation of NO_2_^−^ in the mid-euphotic zone because of physiological and physiochemical differences related to photoinhibition [20] or cell size and redox chemistry [21]. Alternatively, phytoplankton can be subject to multiple factors affecting the degree to which they excrete NO2^−^ during growth on NO_3_^−^ [22]. These factors include light [23], growth rate [24], temperature [25], external NO_3_^−^ concentration [26, 27, 28], Fe limitation [29], and N deficiency [26, 30].

Several features of *Prochlorococcus’* diversity suggest that NO_2_^−^ cycling could be an important facet of this organism’s ecology, particularly for the abundant low-light adapted *Prochlorococcus* that dominate the upper reaches of the nitracline. In N-limited systems, a sizable fraction (20-50%) of *Prochlorococcus* have the capacity for NO_3_^−^ assimilation [10]. Within the low-light adapted LLI clade of *Prochlorococcus*, the NO_2_^−^ assimilation trait appears to be found in most or all cells while the full pathway for NO_3_^−^ assimilation is only observed in a subset of cells [11]. These trait frequencies bear the hallmarks of medium-frequency dependent selection which are often governed by cross-feeding interactions [31]. Given that LLI *Prochlorococcus* live in the vicinity of the primary NO_2_^−^ maximum layer, we hypothesized that those capable of NO_3_^−^ assimilation have the potential for releasing NO_2_^−^ back into the environment – consistent with what is observed in larger size classes of phytoplankton [22]. In this study, we explore incomplete assimilatory NO_3_^−^ reduction by *Prochlorococcus* and further examine the potential for cross-feeding and intra-population cycling of NO_2_^−^, a central intermediate in the N cycle.

## MATERIALS AND METHODS

### Strains

The cultures used in this study included the low-light adapted *Prochlorococcus* strains MIT0915, MIT0917, and MIT1214, the high-light adapted *Prochlorococcus* strain SB, as well as *Synechococcus* strains WH8102 and WH7803 (Table S1). All strains were routinely assayed for heterotrophic contaminants by staining cells with SYBR green and assessing the fluorescence and light scattering properties of both stained and unstained cells using a Guava easyCyte 12HT Flow Cytometer (MilliporeSigma, Burlington, MA, USA) – cultures that did not exhibit the presence of non-photosynthetic cells in the stained samples and had a single cyanobacteria population were presumed axenic and unialgal. All axenic cultures were routinely assessed for purity by confirming a lack of turbidity after inoculation into a panel of purity test broths as described previously [9].

### NO_2_^−^ production rates

The NO_3_^−^ assimilating strains (MIT0915, MIT0917, SB, WH7803, and WH8102) were grown in triplicate as pure cultures using Pro99 medium (natural seawater base; Sargasso Seawater) with the 800 µM ammonium chloride (NH_4_Cl) omitted and replaced by 800 µM sodium nitrate (NaNO_3_). The cultures were grown in 35 mL of medium in borosilicate glass culture tubes at a temperature of 24^°^C and under continuous illumination from white fluorescent lamps at intensities of 6, 36, 52, and 85 µmol photons m^−2^ s^−1^. The LLI strains (MIT0915 and MIT0917) had poor and inconsistent growth at the highest light intensity, so these strains were only examined at the 3 lower light intensities (6, 36, and 52 µmol photons m^−2^ s^−1^). The *Synechococcus* strains (WH7803 and WH8102) were only examined at 36, 52, and 85 µmol photons m^−2^ s^−1^. Cultures were monitored daily by removing 0.5 mL of culture in order to determine cell abundances with flow cytometry and NO_2_^−^ concentrations with the Griess colorimetric method (Supplementary Materials and Methods). Net cell-specific NO_2_^−^ production rates were calculated as the change in NO_2_^−^ concentration relative to the logarithmic mean of cell concentrations – to account for exponential growth – for successive time points during the log-linear portion of the growth curve.

### Experimental populations

MIT1214 was co-cultured, in triplicate, with either MIT0915 or MIT0917 in 35 mL of medium in borosilicate glass culture tubes using NO3^−^ as the sole N source at 24°C and 16 µmol photons m^−2^ s^−1^ of blue light to simulate the conditions under which these cells would coexist in the wild. As controls, all strains were grown as pure cultures, in duplicate, under the same temperature and light conditions. The MIT1214-MIT0915 and MIT1214-MIT0917 co-cultures, as well as the MIT0915 and MIT0917 pure cultures, used Pro99 medium (natural seawater base; Sargasso Seawater) with 800 µM sodium nitrate (NaNO_3_) as the sole N source. Pure cultures of MIT1214 were grown in Pro99 medium using 100 µM sodium nitrite (NaNO_2_) as the sole N source to serve as a control for growth on NO_2_^−^ and in Pro99 medium using 800 µM NO_3_^−^ as the sole N source to serve as control for the absence of growth on NO_3_^−^.

The co-cultures and pure cultures were sampled daily over 2 sequential transfers to monitor cell abundances with flow cytometry, NO_2_^−^ concentrations using the Griess method, and to prepare qPCR filters for strain specific cell abundance measurements (Supplementary Materials and Methods).

### Quantitative PCR assay

Given that the cell size and fluorescence properties of MIT0915, MIT0917, and MIT1214 overlap when examined using flow cytometry (all are LLI clade cells with similar size and chlorophyll content), we used qPCR to assess the cell abundance of each strain in co-culture. For MIT0915 and MIT0917, we used an assay that we had previously developed for detection of *narB* in these strains [10]. For the detection of MIT1214 we designed a qPCR assay to target the *wckA* gene (encoding a polysaccharide pyruvyl transferase family protein) in MIT1214 that is absent in both MIT0915 and MIT0917. Sample processing, standards, reaction conditions, and amplification efficiencies are detailed in the Supplementary Materials and Methods.

## RESULTS AND DISCUSSION

### *Prochlorococcus* strains produce NO_2_^−^ during growth on NO_3_^−^

NO_2_^−^ accumulation and release by phytoplankton growing on NO_3_^−^ can arise from a bottleneck in the NO_3_^−^ assimilation pathway. One possible cause of this bottleneck is limited availability of reducing power – the reduction of NO_3_^−^ requires 2 electrons, while the reduction of NO_2_^−^ requires 6 electrons. Thus, we first asked whether *Prochlorococcus*, as well as the closely-related *Synechococcus*, exhibit incomplete assimilatory NO_3_^−^ reduction with concomitant NO_2_^−^ release [22]. To address this question, we leveraged a collection of strains in the MIT Cyanobacteria Culture Collection (Table S1). One includes the NO_3_^−^ assimilating *Prochlorococcus* SB, a member of the high-light adapted HLII clade of *Prochlorococcus*, which are abundant in warm subtropical surface waters. Among the low-light adapted LLI clade of *Prochlorococcus*, we have identified 3 configurations of the NO_3_^−^ and NO_2_^−^ assimilation gene cassette [11], each represented in our study by the MIT1214, MIT0915, and MIT0917 strains (Table S1). MIT1214 has the capacity for NO_2_^−^ assimilation, but not NO_3_^−^ assimilation – i.e., it has lost the upstream half of the NO_3_^−^ assimilation pathway, but has retained the downstream half for the assimilation of the more reduced NO_2_^−^. MIT0915 can assimilate NO_3_^−^ and also possess a NO_2_^−^ specific transporter (FocA) and a NO_2_^−^ reductase (NirA) that are both closely related to those found in MIT1214. MIT0917 can also assimilate NO_3_^−^, but in contrast to MIT0915, this strain has a divergent version of NirA and has also lost the gene encoding the FocA NO_2_^−^ transporter [11]. The *Synechococcus* strains examined included WH7803 and WH8102. While the environmental distribution of the former is not well resolved, the latter is adapted to warm oligotrophic waters and has a range that overlaps with the abundant HLII clade of *Prochlorococcus* [32]. WH7803 belongs to subcluster 5.1B and has both the FocA NO_2_^−^ transporter and the dual specific NO_3_^−^/NO_2_^−^ NapA transporter. WH8102 belongs to subcluster 5.1A and lacks the FocA NO_2_^−^ transporter.

We looked for evidence of extracellular accumulation of NO_2_^−^ during the growth of *Prochlorococcus* and *Synechococcus* strains on NO_3_^−^ with respect to growth on NH_4_^+^ as the sole N sources. Given that reducing power is ultimately derived from photochemistry in cyanobacteria, we also tested if extracellular accumulation of NO_2_^−^ might be enhanced during growth at lower light intensities. We found that *Prochlorococcus* MIT0917 cultures accumulated substantial concentrations of NO2^−^ when provided NO_3_^−^ as the sole N source (Fig. 1), compared to growth on NH_4_^+^ (Fig. S1). In contrast, NO_2_^−^ concentrations remained below 1 µM (the lower limit of the dynamic range for our assay) in cultures of the closely-related MIT0915 strain during growth on NO_3_^−^ (Fig. 1). The high-light adapted *Prochlorococcus* SB did accumulate NO_2_^−^ in the culture medium during growth on NO_3_^−^, but at substantially lower concentrations compared to MIT0917 (Fig. 1). In comparison to the *Prochlorococcus* strains examined, neither of the *Synechococcus* strains produced levels of NO_2_^−^ that exceeded 1 µM during growth on NO_3_^−^ or NH_4_^+^ (Figs. 2 and S2). Overall, these data show that some strains of *Prochlorococcus* exhibit incomplete assimilatory NO_3_^−^ reduction with concomitant release of NO_2_^−^.

**Fig. 1.**
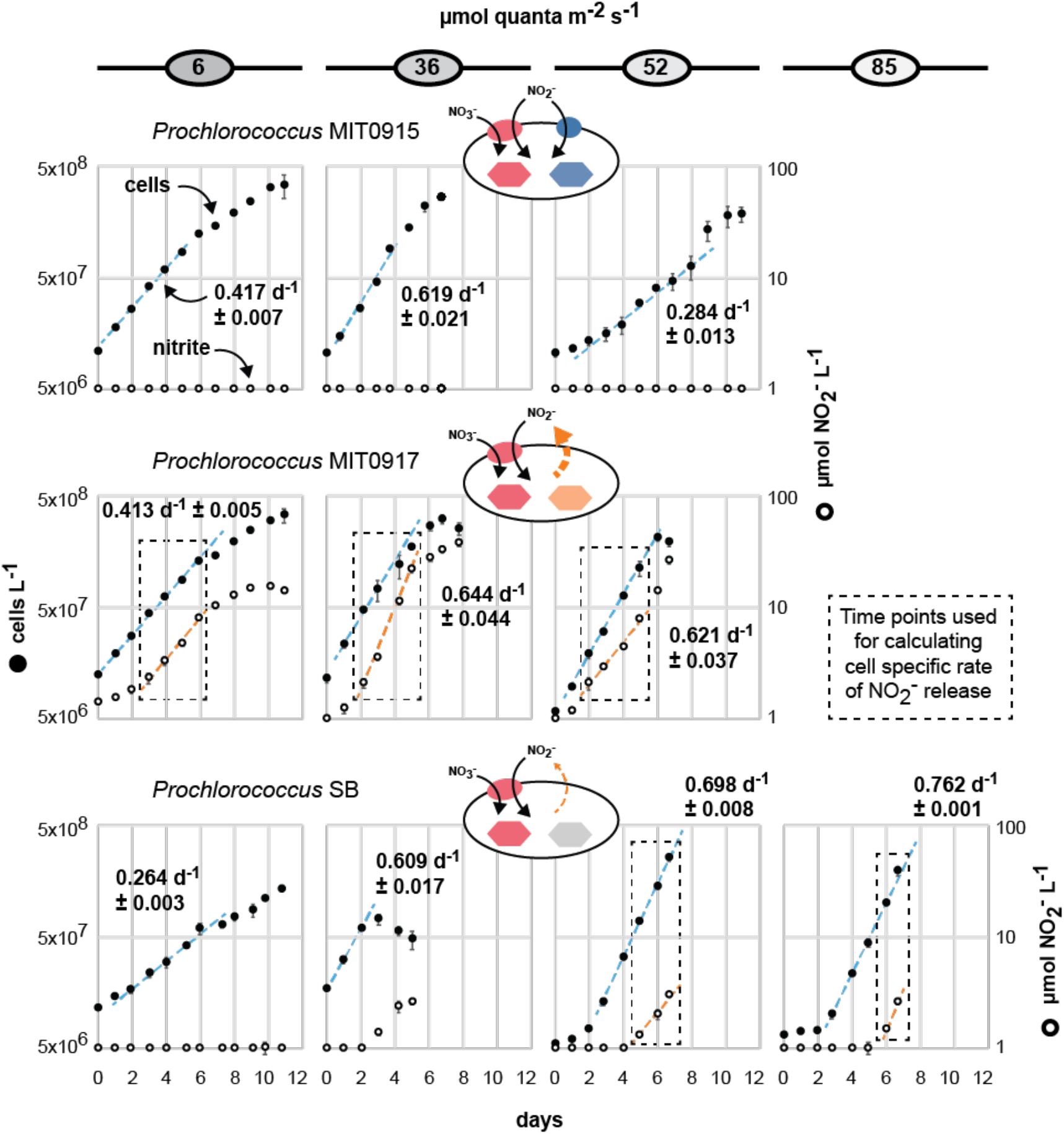
NO_2_^−^ accumulation in batch cultures of *Prochlorococcus* during growth on NO_3_^−^ as the sole N source over a range of light intensities. Both *Prochlorococcus* MIT0917 (LLI clade) and *Prochlorococcus* SB (HLII clade) were observed to accumulate extracellular NO_2_^−^ when provided NO_3_^−^ as the sole N source. NO_2_^−^ concentrations below the dynamic range of the assay (< 1 µM NO_2_^−^) are plotted on the x-axis. Cell-specific rates of NO_2_^−^ release (nmol NO_2_^−^ cell^−1^ d^−1^) were calculated from the log-linear portion of the growth curve.

**Fig. 2.**
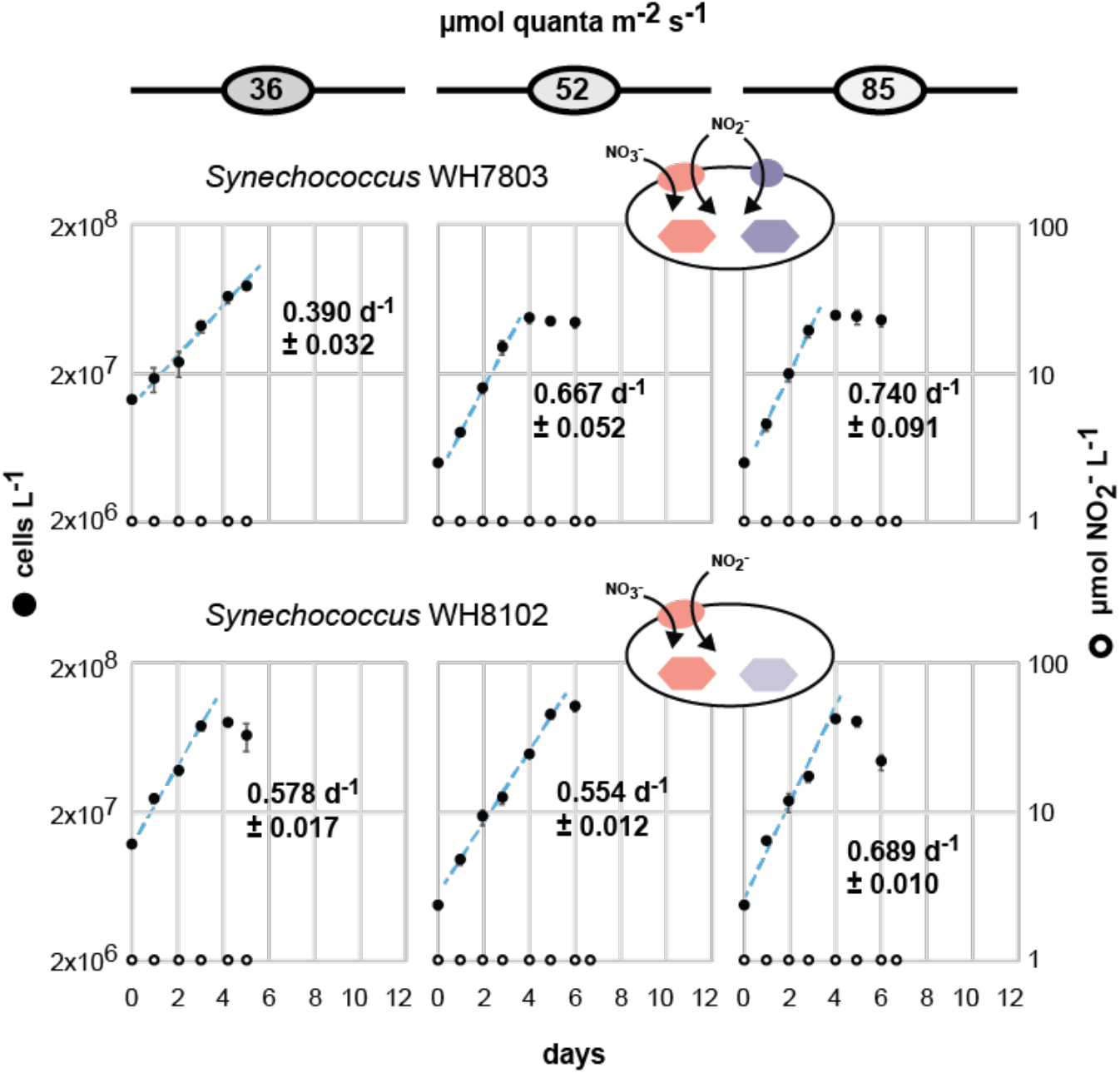
NO_2_^−^ accumulation in batch cultures of *Synechococcus* during growth on NO_3_^−^ as the sole N source over a range of light intensities. Neither *Synechococcus* strain accumulated extracellular NO_2_^−^. NO_2_^−^ concentrations below the dynamic range of the assay (< 1 µM NO_2_^−^) are plotted on the x-axis.

We next examined the net cell-specific NO_2_^−^ production rates of MIT0917 and SB during growth on NO_3_^−^. The rates of NO_2_^−^ excretion by the low-light adapted *Prochlorococcus* MIT0917 was significantly greater than that for the high-light adapted *Prochlorococcus* SB (Fig. 3). At the same light intensity of 52 µmol photons m^−2^ s^−1^, MIT0917 produced NO_2_^−^ at a 5-fold higher rate compared to SB (Fig. 3). The rate of NO_2_^−^ production by MIT0917 was highest at 36 and 52 µmol photons m^−2^ s^−1^ in comparison to the lowest light intensity examined. These rates are necessarily inclusive of potential reuptake of NO_2_^−^ by the dual specific NO_3_^−^/NO_2_^−^ NapA transporter encoded by both of these strains. Nevertheless, reuptake of NO_2_^−^ could be limited by the fact that the molar equivalents of NO_3_^−^ would exceed that of NO_2_^−^ by at least one order of magnitude throughout the entire growth curve.

**Fig. 3.**
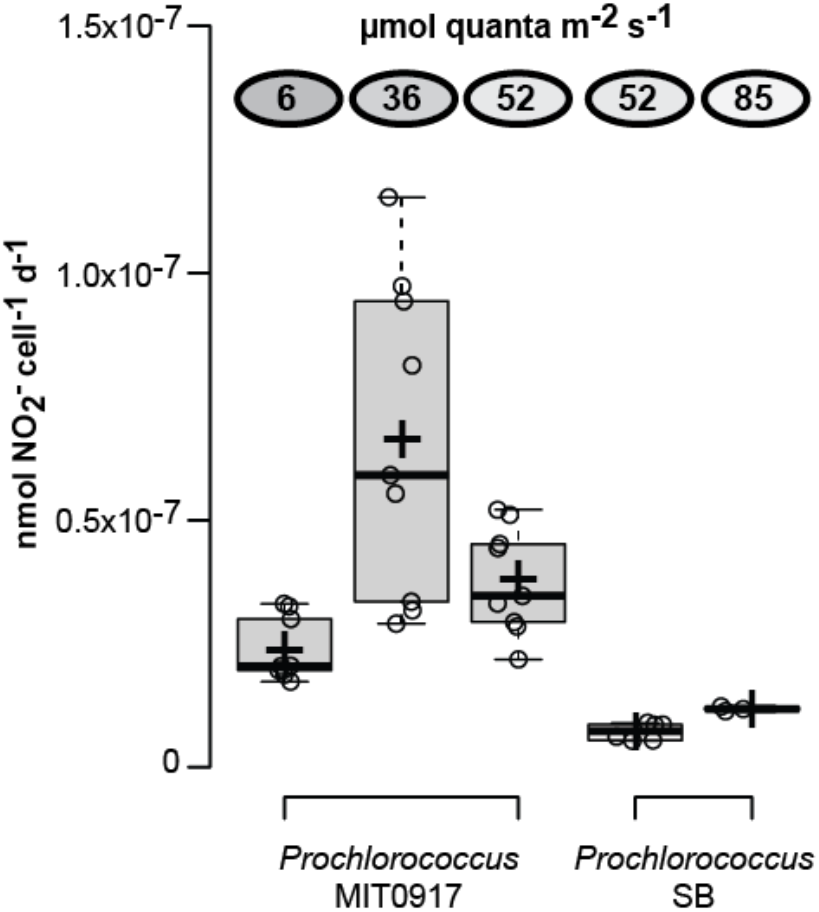
Cell-specific NO_2_^−^ production rates for two strains of *Prochlorococcus*, as a function of light intensity, when grown on NO_3_^−^ as the sole N source. Mean (+/- standard deviation) of NO_2_^−^ production rates for MIT0917 were 2.4 × 10^−8^ (+/- 6.4 × 10^−9^), 6.6 × 10^−8^ (+/- 3.2 × 10^−8^), and 3.8 × 10^−8^ (+/- 1.1 × 10^−8^) nmol NO_2_^−^ cell^−1^ d^−1^ at light intensities of 6, 36, and 52 µmol photons m^−2^ s^−1^, respectively. SB produced nitrite at rates of 7.2 × 10^−9^ (+/- 1.7 × 10^−9^) and 1.2 × 10^−8^ (+/- 5.0 × 10^−10^) nmol NO_2_^−^ cell^−1^ d^−1^ at light intensities of 52 and 85 µmol photons m^−2^ s^−1^, respectively. Medians are denoted by solid black lines and means are denoted by crosses.

We next wondered how consequential these rates of NO_2_^−^ release by MIT0917 were with respect to the N requirements of low-light adapted *Prochlorococcus*. To evaluate this question, we assumed a cellular N quota of 4.3 fg N cell^−1^ for LLI *Prochlorococcus* [33] and minimal reuptake of NO_2_^−^. Based on these assumptions, we find that approximately 20-30% of the NO_3_^−^ transported into the cell by MIT0917 during exponential growth was released as NO_2_^−^, with the balance assimilated into biomass (i.e., the proportion of N as extracellular NO_2_^−^ relative to the combined N in both biomass and extracellular NO_2_^−^). Overall, these data demonstrate that some strains belonging to an abundant low-light adapted clade of *Prochlorococcus* can release high amounts of NO_2_^−^ when provided NO_3_^−^ as the sole N source.

### Contrasting features of NO_3_^−^ and NO_2_^−^ flow in *Prochlorococcus* and *Synechococcus*

Based on what is known about the gene content [11] and physiology (Figs. 1 and 2) of these strains, we can assign hypothetical pathways of inorganic N flow (Fig. 4). For LLI *Prochlorococcus*, these hypothetical pathways map onto the 3 distinct configurations of the NO_3_^−^ and NO_2_^−^ assimilation gene cassette that we have observed in their genomes [11]. Many LLI *Prochlorococcus* have the configuration represented by MIT1214 – that is, they can assimilate both NH_4_^+^ and NO_2_^−^, but not NO_3_^−^. The remaining LLI *Prochlorococcus* – those represented by the NO_3_^−^ assimilating strains, MIT0915 and MIT0917 – have contrasting features with regard to NO_2_^−^ production potential (Fig. 4). Under the N-replete conditions we examined, MIT0917 accumulated high extracellular quantities of NO_2_^−^ relative to the proportion of N that was ultimately assimilated into biomass. The closely related MIT0915 strain, however, released negligible quantities of NO_2_^−^ under the same conditions.

**Fig. 4.**
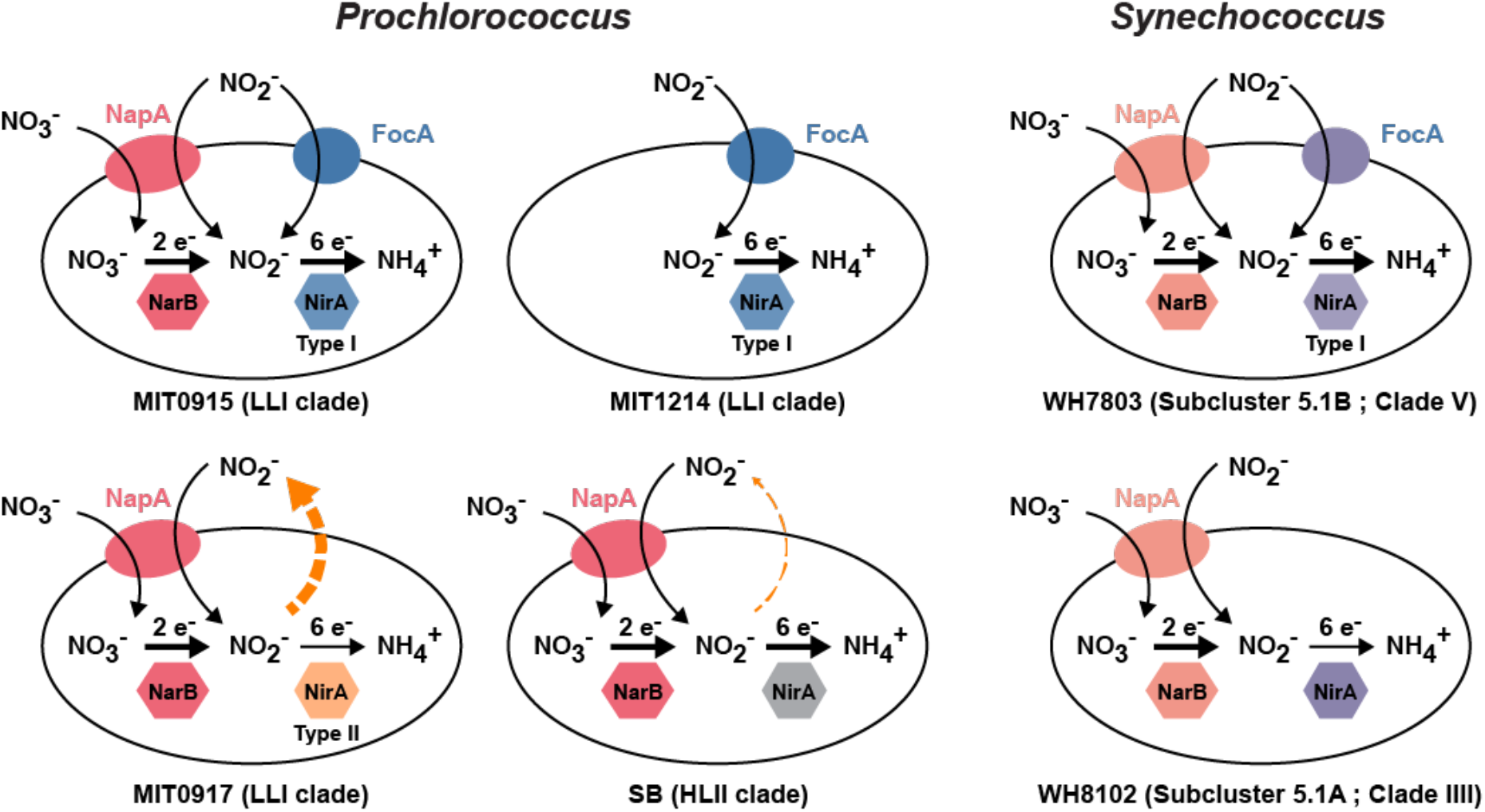
Hypothetical cellular inorganic N inputs and outputs for *Prochlorococcus* and *Synechococcus* strains are based on physiology data (Figs. 1 and 2) and functional annotation of pathway proteins [11]. SB lacks a dedicated NO_2_^−^ transporter (FocA), but can take up NO_2_^−^ using the dual-specific NO_3_^−^/NO2^−^ NapA transporter. MIT0917 has a Type II NirA and lacks the FocA NO_2_^−^ transporter. Both SB and MIT0917 excrete NO_2_^−^ (dashed orange arrow). MIT0915 has a Type I NirA and has also retained the FocA NO_2_^−^ transporter – MIT0915 did not excrete NO_2_^−^ during growth on NO_3_^−^ (Fig. 1). MIT1214 has lost the upstream half of the NO_3_^−^ assimilation pathway, but has retained FocA and the Type I NirA for the transport and assimilation of NO_2_^−^. In comparison to the *Prochlorococcus* strains examined, neither *Synechococcus* strain releases NO_2_^−^ during growth on NO_3_^−^. WH7803 possesses the FocA NO_2_^−^ transporter, while WH8102 does not.

*Prochlorococcus* SB produced much less NO_2_^−^ during growth on NO_3_^−^ than *Prochlorococcus* MIT0917 – and only at the highest light intensities examined – but, similar to MIT0917, SB also lacks the FocA NO_2_^−^ transporter (Fig. 4). *Prochlorococcus* SB belongs to the high-light adapted HLII clade, which is the most abundant clade of *Prochlorococcus* globally and can represent >90% of depth integrated *Prochlorococcus* in warm tropical and subtropical waters [15]. Although the culturing conditions we employed are quite different than those that cells experience in the wild, HLII clade cells with the potential for NO_2_^−^ release – even if less than that of some LLI clade cells – could have an important impact on NO_2_^−^ cycling in the global ocean due to their sheer abundance. *Synechococcus*, while broadly distributed and responsible for a greater fraction of net primary production compared to *Prochlorococcus*, does not appear to release NO_2_^−^ in batch culture (Figs. 2 and 4). An important caveat is that the strains we examined represent only a fraction of the diversity of *Synechococcus*.

### Wild populations of LLI clade *Prochlorococcus* are composed of coexisting functional types delineated by their use of NO_3_^−^ and NO_2_^−^

In both the subtropical North Pacific and North Atlantic oceans, LLI *Prochlorococcus* (either with or without the capacity for NO_3_^−^ assimilation) often reach maximum abundance within the subsurface chlorophyll maximum layer [15] and in the vicinity of the primary NO_2_^−^ maximum layer [10]. Our data now indicate that there is a significant degree of phenotypic diversity with respect to the use of NO_2_^−^ and NO_3_^−^ and that this functional diversity maps onto the genomic diversity of the LLI clade of *Prochlorococcus*. Given these observations, how are LLI *Prochlorococcus* populations structured with respect to the 3 distinct functional types that we have identified (Fig. 4)? To address this question, we turned to metagenomic data [34] from the subsurface chlorophyll maximum layer at time-series stations in the North Pacific Subtropical Gyre (Hawai’i Ocean Time-series; HOT) and the North Atlantic Subtropical Gyre (Bermuda Atlantic Time-series Study; BATS).

In both the North Pacific and the North Atlantic subtropical gyres, we observed that MIT1214-like cells (those that assimilate NO_2_^−^ but not NO_3_^−^) dominated the LLI *Prochlorococcus* populations throughout the year and generally exceeded 60% of total LLI genomes (Fig. 5). In the North Pacific ecosystem, overall frequencies of each functional type were quite stable on seasonal time scales, with each of the NO_3_^−^ assimilating functional types making up roughly 15% of the population (Fig. 5). In contrast, the seasonal dynamics in the North Atlantic ecosystem were readily apparent. Frequencies of the NO_3_^−^ assimilating functional types waned in the winter and spring and then increased through the summer and autumn (Fig. 5). These observations are consistent with previous work that demonstrated higher abundances of HLII *Prochlorococcus* with the potential for NO_3_^−^ assimilation during the summer and autumn in the North Atlantic [10], when overall surface N concentrations were low. The MIT0915-like functional type (the one that did not exhibit incomplete assimilatory NO_3_^−^ reduction in our culture experiments) generally dominated the NO_3_^−^ assimilating functional types in the North Atlantic ecosystem’s LLI *Prochlorococcus* populations (Fig. 5). In contrast, higher frequencies of the MIT0917-like functional type (the one that releases NO_2_^−^) in the North Pacific compared to the North Atlantic suggests its configuration of the NO_3_^−^ assimilation gene cassette may be better adapted to the stable, well-stratified, and generally N-limited waters of the North Pacific Subtropical Gyre.

**Fig. 5.**
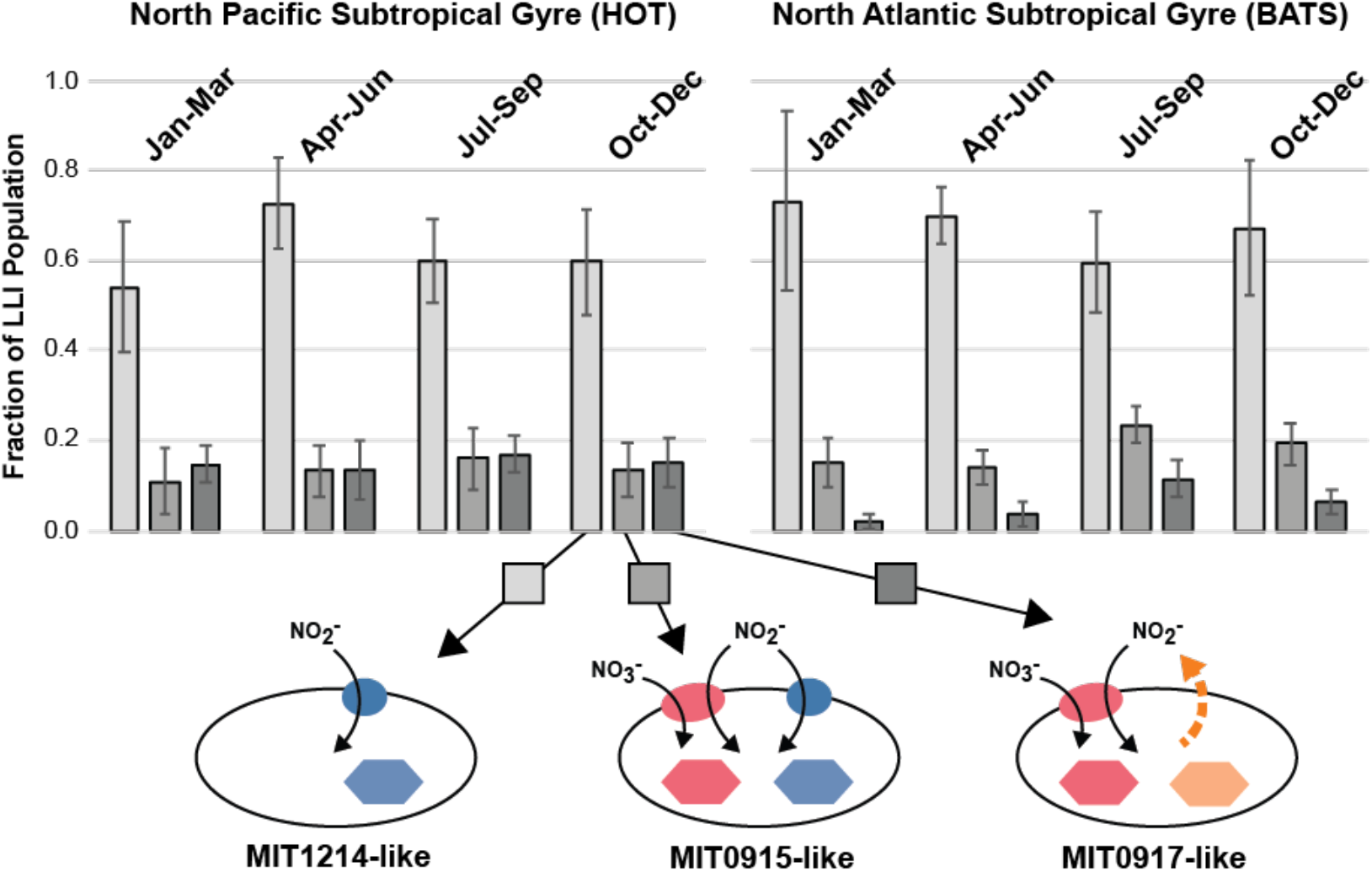
Distribution of functional types in the LLI clade *Prochlorococcus* (based on N flow pathways in Fig. 4) in the North Pacific and North Atlantic subtropical gyres. NO_3_^−^ assimilating genotypes are abundant in the North Pacific throughout the year. Putative NO_2_^−^ producers (MIT0917-like) reach maximum abundance in the North Atlantic during summer stratification of the water column. NO_2_^−^ consumers (MIT1214-like) typically account for >50% of the LLI populations in both ecosystems.

### *Prochlorococcus* exchange N in experimental populations

Given that LLI *Prochlorococcus* with different N assimilation features (Fig. 4) coexist in the marine environment (Fig. 5), we next assessed whether strains with different N assimilation genotypes could form metabolic dependencies in the laboratory. To achieve this, we co-cultured *Prochlorococcus* MIT1214 (which can use NO_2_^−^ but not NO_3_^−^) with either *Prochlorococcus* MIT0915 or *Prochlorococcus* MIT0917 (each of which can use both NO_3_^−^ and NO_2_^−^) in medium containing NO_3_^−^ as the sole N source.

Relative to pure cultures of MIT0917 (Fig. 6B, F), co-cultures of MIT1214 and MIT0917 did not accumulate NO_2_^−^ in the culture medium during balanced exponential growth (Fig. 7A, B), because any NO_2_^−^ released by MIT0917 was used to fulfill the N requirements of MIT1214.

**Fig. 6.**
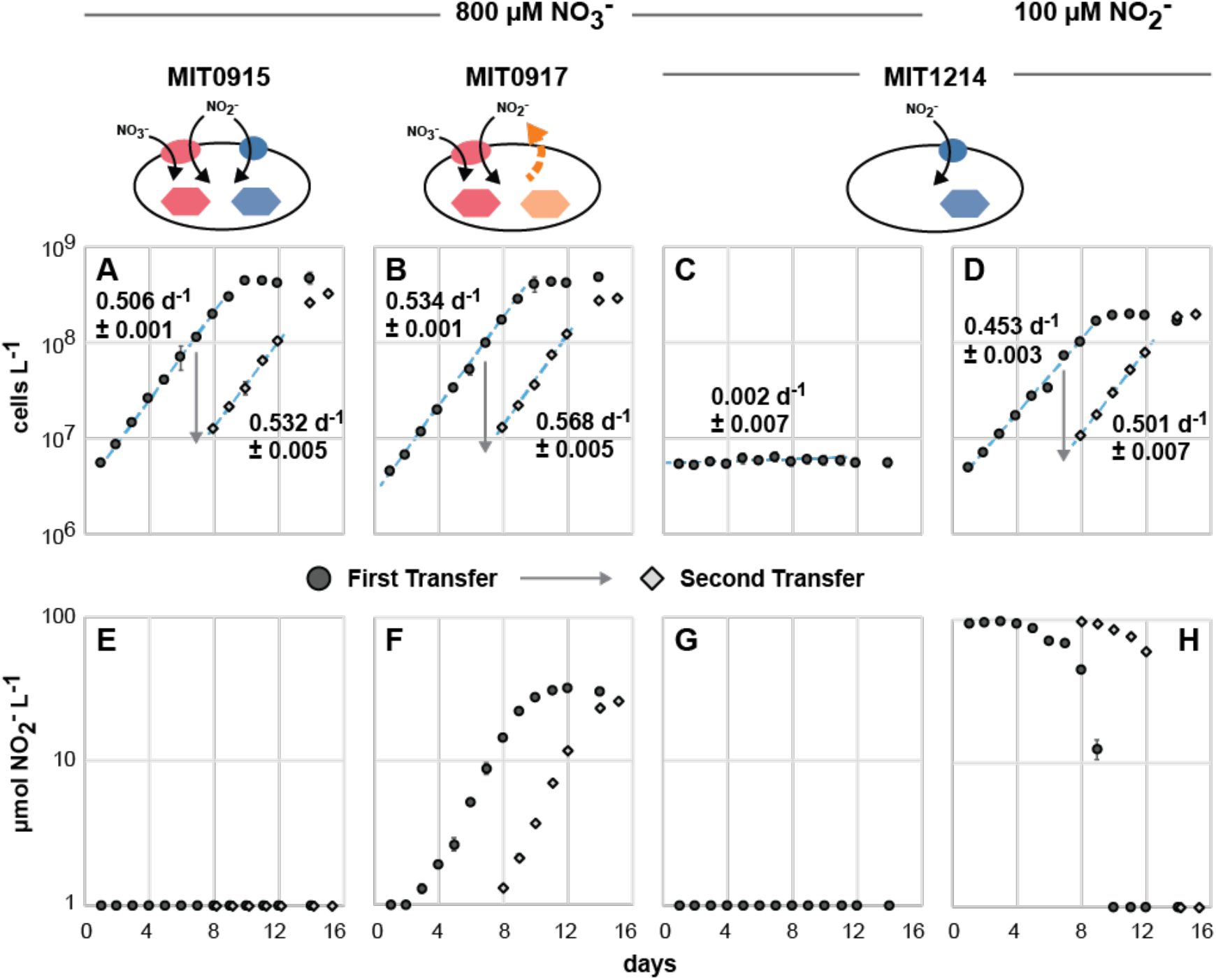
Pure cultures of MIT0915, MIT0917, and MIT1214 followed over 2 transfers (black circles followed by gray diamonds) as contemporaneous controls for co-culture experiments. As pure cultures, MIT0915 and MIT0917 grow using NO_3_^−^ (A, B), but only MIT0917 does so with concomitant release of NO_2_^−^ (E, F). MIT1214 cannot use NO_3_^−^ as a N source (C), but has retained the genes necessary for growth on NO_2_^−^ (D, H).

**Fig. 7.**
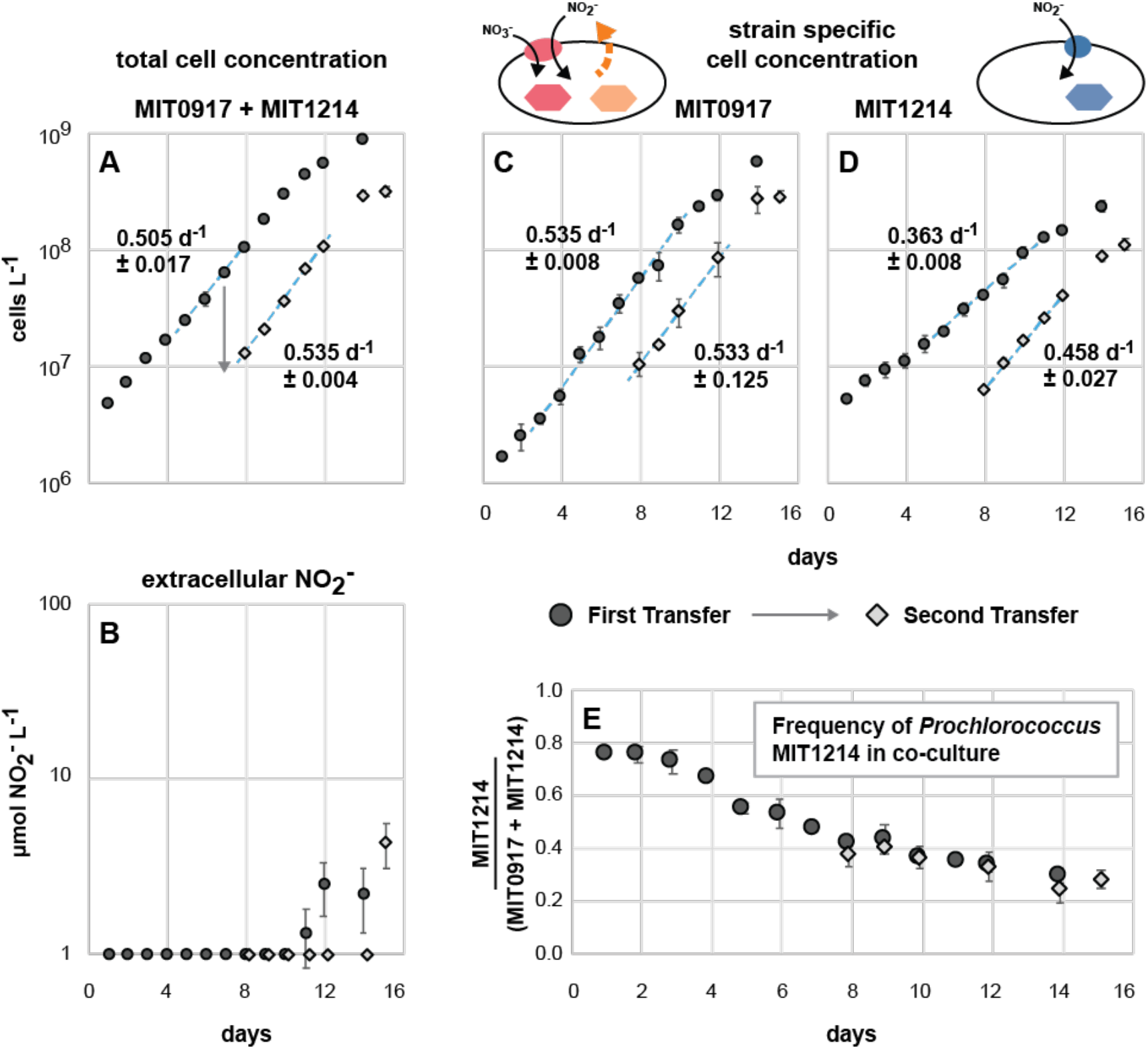
Co-culture of *Prochlorococcus* MIT0917 (NO_2_^−^ producer) and *Prochlorococcus* MIT1214 (NO_2_^−^ consumer). Total cell counts and specific growth rates were determined by flow cytometry (A). NO_2_^−^ does not accumulate in the co-culture during balanced exponential growth (B), instead being drawn down by MIT1214. The growth rates of the individual strains in the co-culture, assessed by quantitative PCR (C, D), were similar to those growing as pure cultures (Fig. 6). The frequency of MIT1214 in the co-culture converged at approximately 30% of total cell numbers after 2 transfers (E).

Some NO_2_^−^ accumulation was observed as the MIT1214-MIT0917 co-culture approached stationary phase (Fig. 7A, B), suggesting an imbalance between production and consumption of NO_2_^−^ outside of balanced exponential growth. The growth rates of each of the two strains in co-culture (Fig. 7C, D) were similar to their growth rates in pure culture (Fig. 6B, D), suggesting that MIT0917 could supply nearly all of the N needs of MIT1214. Further, we expect that the growth of MIT1214 under these conditions resembles growth in continuous culture systems – i.e., in co-culture, MIT1214 was likely poised at some degree of N limitation with growth controlled by the rate of NO_2_^−^ release by the partner strain. In these co-cultures, the frequency of the MIT1214 strain settled at roughly 30% of total cell numbers as determined by quantitative PCR (Fig. 7E) – providing additional support for our estimate that MIT0917 partially reduces and excretes up to 30% of the NO_3_^−^ transported into the cell as extracellular NO_2_^−^.

When MIT1214 was paired with MIT0915, a strain that does not produce NO_2_^−^ when growing on NO_3_^−^ in pure culture (Fig. 6A, E), we expected the growth of MIT1214 to stop after any carry-over NO_2_^−^ from the inoculum was exhausted. On the contrary, MIT1214 continued to grow (Fig. 8D), but at significantly lower growth rates compared to either growth in the presence of MIT0917 using NO_3_^−^ as the N source (Fig. 7D) or to the growth of MIT1214 in pure culture using NO_2_^−^ (Fig. 6D). As expected, NO_2_^−^ did not accumulate during the growth of the MIT1214-MIT0915 co-culture (Fig. 8A, B). The frequency of the MIT1214 strain was driven to <5% of total cell numbers over the course of 2 sequential transfers (Fig. 8E), likely as a consequence of the MIT1214 strain’s depressed growth rate in the presence of the MIT0915 strain. The continued slow growth of MIT1214 suggests the possibility that MIT0915 does produce low quantities of NO_2_^−^ when growing on NO_3_^−^. Recall that MIT0915 also possesses a FocA NO_2_^−^ specific transporter (Fig. 4) – in contrast to MIT0917 – which may facilitate some reuptake of NO_2_^−^ and thus maintain undetectable NO_2_^−^ concentrations in pure culture (Fig. 6A, E). Supply of NO_2_^−^ at rates that keep the concentration of NO_2_^−^ at or below the half-saturation constant (Ks) for MIT1214’s growth on NO_2_^−^ would be consistent with its significantly lower growth rate when co-cultured with MIT0915.

**Fig. 8.**
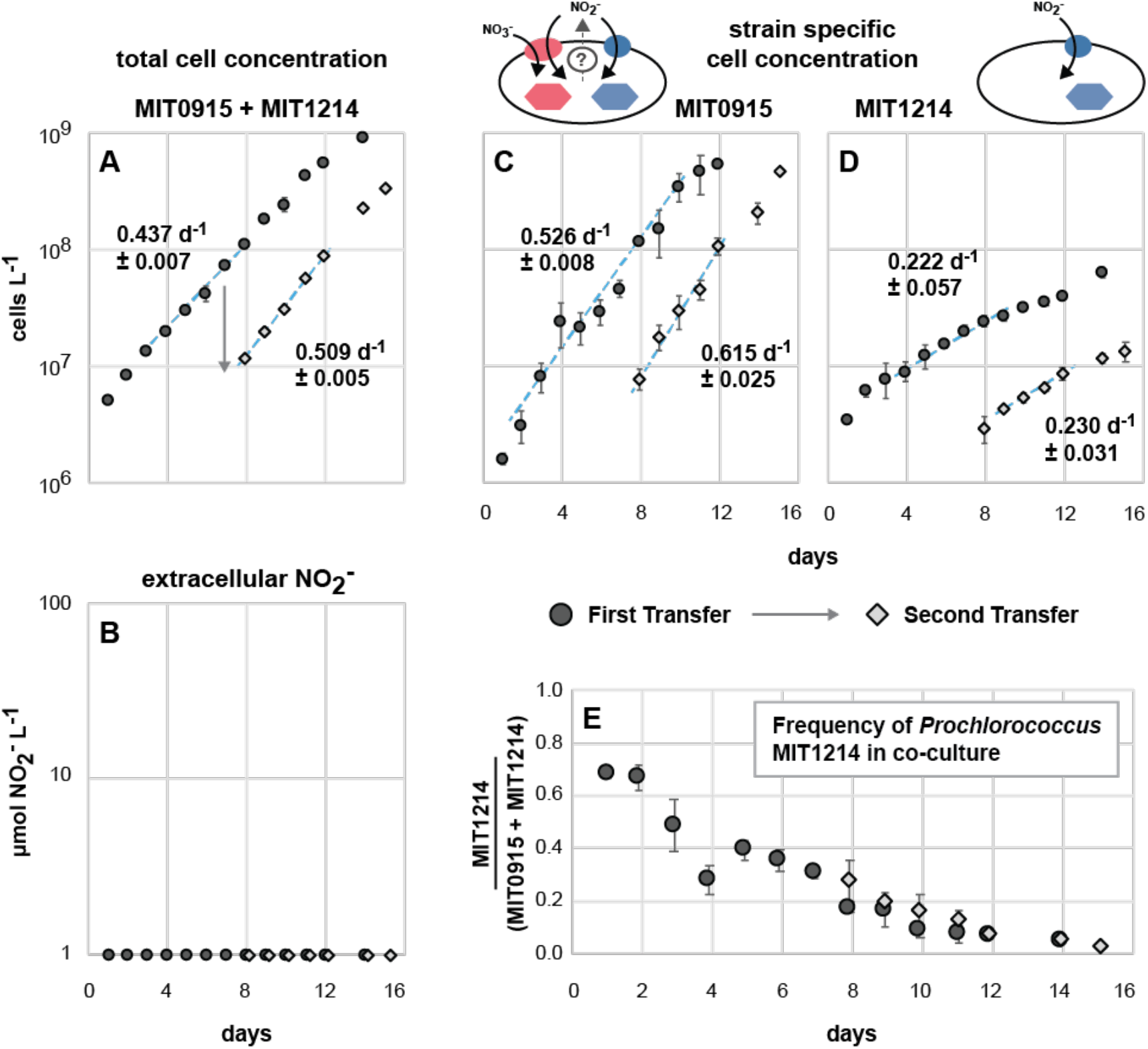
Co-culture of *Prochlorococcus* MIT0915 and *Prochlorococcus* MIT1214. Total cell counts and specific growth rates were determined by flow cytometry (A). NO_2_^−^ does not accumulate in the co-culture (B) as observed with pure cultures of MIT0915 (Fig. 6). As determined by quantitative PCR, MIT0915 maintained steady growth (C) while the growth rate of MIT1214 (D) declined by more than 50% compared to growth as a pure culture using NO_2_^−^ as the N source (Fig. 6). Although MIT1214 was driven to less than 5% of total cell numbers after 2 transfers (E), continued growth of the strain at a depressed growth rate suggests some level of cryptic NO_2_^−^ release by the MIT0915 strain (denoted in the cartoon schematic of N flow in MIT0915).

### Causes and consequences of incomplete assimilatory NO_3_^−^ reduction by *Prochlorococcus*

Our work has uncovered new potential links between *Prochlorococcus* and the N cycle by demonstrating that some *Prochlorococcus* divert a sizable fraction of transported NO3^−^ to extracellular pools of NO_2_^−^. An intriguing feature of this phenomenon is the degree of phenotypic variability between strains. For *Prochlorococcus* cultures, the ability of some to accumulate NO_2_^−^ (Figs. 1, 3, and 6F) maps onto each strain’s particular version of the NO_3_^−^ assimilation gene cluster [11]. Given the global abundance of *Prochlorococcus* in the world’s oceans, the fraction of *Prochlorococcus* that release NO_2_^−^ when reducing NO_3_^−^ could have important consequences for our understanding of primary production and N cycle processes that occur in the tropical and subtropical ocean.

While many eukaryotic phytoplankton exhibit incomplete assimilatory NO_3_^−^ reduction, this process is not well constrained for the picocyanobacteria that dominate the expansive subtropical gyres of the open ocean. Why would an organism such as *Prochlorococcus*, which is well-adapted to life in oligotrophic habitats, release N back into its environment when this nutrient is often in limited supply? Our observations could be related to laboratory culture conditions where the concentrations of inorganic nutrients are much higher than would be observed in the field. Yet, only one functional type of LLI *Prochlorococcus* (represented by the MIT0917 strain) exhibited incomplete assimilatory NO_3_^−^ reduction. It is possible that the divergent version of the NirA NO_2_^−^ reductase possessed by MIT0917 – and similar *Prochlorococcus* in the field – has distinct biochemical features. One hypothesis is that this divergent NirA has a higher substrate affinity, perhaps providing these cells with an advantage under chronically N limited conditions. Under replete conditions in batch culture, these cells might experience a kinetic bottleneck at the NO_2_^−^ reduction step of the NO_3_^−^ assimilation pathway that results in cellular accumulation of NO_2_^−^ because NO_3_^−^ reduction outpaces the k_cat_ of this divergent NirA. This NO_2_^−^ could then diffuse out of the cell as nitrous acid (HNO_2_) which would make up a small fraction of the intracellular NO_2_^−^ pool at cellular pH [35]. Analogous field conditions would be intermittent upwellings of NO_3_^−^ that temporarily increase local substrate supply [36].

*Prochlorococcus* MIT0917 also lacks the FocA NO2^−^ specific transporter that is found in other LLI *Prochlorococcus* (Fig. 4), including the MIT0915 strain that does not exhibit extracellular accumulation of NO_2_^−^ in batch culture. An alternative, but not mutually exclusive, hypothesis for NO_2_^−^ release by MIT0917 is that this strain lacks the capacity for cyclic retention of NO_2_^−^ that may diffuse out of the cell as HNO_2_. It is common for bacteria to employ so-called “futile cycles” to mitigate the loss of nutrients and metabolites across the cell membrane [37, 38]. While this process expends energy, it can serve to regulate substrate retention and maintain sufficiently high concentrations of a substrate within the cell. MIT0917 may not require re-uptake of NO_2_^−^ if its NirA is optimized for low internal substrate concentrations – in such a scenario, the *focA* gene could have been lost due to a general bias towards gene deletion in the absence of a sufficient selective advantage. The MIT0915 strain, on the other hand, might require a mechanism for cyclic retention of NO_2_^−^ if its NirA has a lower substrate affinity. Notably, we did observe slow growth of MIT1214 when in co-culture with MIT0915 (Fig. 8D), suggesting that MIT0915 might release some quantities of NO_2_^−^ to the extracellular milieu – this would put it in competition with MIT1214 for access to NO_2_^−^ in order to maintain sufficiently high cellular NO_2_^−^ concentrations.

Our laboratory-based observations suggest that *Prochlorococcus* may interact with the primary NO_2_^−^ maximum layer in complicated ways. Although this feature is ubiquitous in stratified oceanic water columns, it is still unclear what mechanisms are responsible for its emergence. In addition to eroding the primary NO_2_^−^ maximum layer through NO_2_^−^ uptake and assimilation, *Prochlorococcus* also appears to have the potential to amplify the magnitude of the primary NO_2_^−^ maximum layer. At present, it is unclear if *Prochlorococcus* exhibit NO_2_^−^ release in the wild and, if so, what abiotic and biotic factors might influence NO_2_^−^ cycling in these populations.

Extrapolating from NO_2_^−^ production rates for both *Prochlorococcus* and ammonia-oxidizing archaea as well as the abundance of marker genes for these microbes in the field, we postulate that NO_2_^−^ production by *Prochlorococcus* could be responsible for some degree of NO_2_^−^ produced in the euphotic zone. *Prochlorococcus* MIT0917 releases NO_2_^−^ at rates from 2 × 10^−8^ nmol NO_2_^−^ cell^−1^ d^−1^ up to 7 × 10^−8^ nmol NO_2_^−^ cell^−1^ d^−1^ (Fig. 3). In comparison, the dominant ammonia-oxidizing microorganism in subtropical open ocean ecosystems, *Candidatus* Nitrosopelagicus brevis [39], produces NO_2_^−^ at a rate of about 2 × 10^−6^ nmol NO_2_^−^ cell^−1^ d^−1^ in batch culture [40]. Ammonia-oxidizing archaea are thus expected to produce NO_2_^−^ at 30-100 fold higher rates than *Prochlorococcus*, but the latter is often more abundant. For instance, ammonia-oxidizing archaea in the epipelagic zone have been typically observed at abundances of 1,000-6,000 *amoA* gene copies mL^−1^ [21, 41, 42]. Low-light adapted *Prochlorococcus* with the capacity of NO_3_^−^ assimilation, however, can reach abundances of 1,000-40,000 cells mL^−1^ in the vicinity of the subsurface chlorophyll maximum layer [10]. Therefore, depending on both rates and cell abundances, *Prochlorococcus* could be responsible from anywhere between <1% to approximately 50% of NO_2_^−^ production in the mid-euphotic zone as a fraction of *Prochlorococcus* and nitrifier derived NO_2_^−^. Under some conditions, *Prochlorococcus* might rival the NO_2_^−^ production by ammonia-oxidizing archaea.

As we have demonstrated, different functional types of *Prochlorococcus* can coexist under conditions where NO_2_^−^ cross-feeding is promoted and NO_2_^−^ accumulation is minimized (Fig. 7). Consequently, NO_2_^−^ production and consumption in *Prochlorococcus* populations might be a cryptic process whereby there is no net accumulation of NO_2_^−^ at steady-state. In the wild, *Prochlorococcus* populations could dynamically assemble in response to the availability of N sources of varying redox state as well as in response to community-wide competition for these N sources. Net NO_2_^−^ accumulation might only occur within these populations during periods of perturbation (e.g., changes in light intensity or nutrient supply). Additional study is warranted to examine the conditions under which *Prochlorococcus* populations are either net producers or net consumers of NO_2_^−^ and evaluate how microbial populations and communities modulate the availability of various N sources that ultimately impact production and remineralization processes in N-limited systems. At the population level, the dynamic assembly of distinct functional types of *Prochlorococcus* could emerge through interactions that are mediated, in part, by cross-feeding of NO2^−^. We posit that trait variability and the selection of complementary functions might facilitate robustness or resiliency in microbial populations. *Prochlorococcus*, as a key primary producer in the tropical and subtropical ocean, offers an extremely valuable lens through which to constrain the rules under which emergent features arise.

## ACKNOWLEDGEMENTS

This work was supported by grants from the National Science Foundation (OCE-2048470 to P.M.B.) and the Simons Foundation (Life Sciences Project Award ID 337262, S.W.C; SCOPE Award ID 329108, S.W.C.). The authors thank Rogier Braakman (MIT) for insightful discussions. This paper is a contribution from the Simons Collaboration on Ocean Processes and Ecology (SCOPE).

## COMPETING INTERESTS

The authors declare no competing financial interests.

## Notes

### Competing Interest Statement

The authors have declared no competing interest.

