## Supplementary Appendix for "Production and cross-feeding of nitrite within *Prochlorococcus* populations"

**This PDF file includes:**

- Supplementary Materials and Methods
- Table S1 – *Prochlorococcus* and *Synechococcus* strains
- Fig. S1 – Extracellular $NO_2^-$ during the growth of *Prochlorococcus* on $NH_4^+$
- Fig. S2 – Extracellular $NO_2^-$ during the growth of *Synechococcus* on $NH_4^+$
- Supplementary References

SUPPLEMENTARY MATERIALS AND METHODS

**Strains.** The cultures used in this study are detailed in Table S1. *Prochlorococcus* MIT1214 was isolated from an enrichment culture initiated as part of the Hawaii Ocean Experiment-Dynamics of Light and Nutrients (HOE-DYLAN) VII expedition (UNOLS Cruise ID: KM1217) at coordinates 22.8$^o$N and -158.0$^o$E. Unfiltered seawater from 175m was aliquoted into acid-washed 30 mL polycarbonate oakridge tubes and amended with 16 µM sodium nitrate, 1 µM sodium phosphate, 0.117 µM ethylenediaminetetraacetic acid, 0.117 µM iron(III) chloride, 0.0090 µM manganese(II) chloride, 0.0008 µM zinc(II) sulfate, 0.0005 µM cobalt(II) chloride, 0.0003 µM sodium molybdate, 0.0010 µM sodium selenite, and 0.0010 µM nickel(II) chloride. Enrichment cultures were routinely monitored by flow cytometry and those exhibiting growth of a *Prochlorococcus*-like population were transferred into fresh medium and ultimately acclimated to growth on Pro99 medium. As part of our study, MIT1214, MIT0915, and MIT0917 were rendered axenic by dilution to extinction [1].

**$NO_2^-$ determination.** Extracellular $NO_2^-$ concentrations were determined via the Greiss colorimetric method that uses sequential additions of sulfanilamide and *N*-(1-naphthyl)ethylenediamine (NED) to produce a pink-red azo dye with a maximum absorption at a wavelength of 540 nm. A solution of 1% (10 mg $mL^{-1}$) sulfanilamide, 5% 12M HCl, and 0.1% (1 mg $mL^{-1}$) NED was filtered through a 0.2um filter into UV resistant bottles. Aliquots of a 1 mM sodium nitrite ($NaNO_2$) standard solution was stored frozen at -20$^o$C and thawed daily to prepare dilutions spanning 1-50 µM for the generation of a standard curve. To prepare samples for quantification of $NO_2^-$ , 0.15 mL was removed from cultures and filtered through a 96-well 0.45µm MultiScreenHTS HVfilter plate (MilliporeSigma, Burlington, MA, USA) capable of capturing >99% of *Prochlorococcus* cells. Dilutions of the $NaNO_2$ standard were filtered in the same plate as the culture samples to ensure similar treatment. 100uL of filtrate was then transferred from each well to a flat-bottomed, 96-well microplate. Sulfanilamide (50 uL) was added to each well, mixed by pipetting, and incubated in the dark for 10 minutes to allow for chromophore formation. Subsequently, NED (50 uL) was added to each well, mixed by pipetting, and incubated in the dark for an additional 10 minutes to allow for coupling and color development. Absorbance at 540 nm was then determined by using a Synergy 2 Plate Reader (BioTek Instruments, Winooski, VT, USA).

**Setup and sampling of experimental populations.** Pure cultures of MIT0915, MIT0917, and MIT1214 were passaged twice at 24ºC and 16 µmol photons $m^{-2}$ $s^{-1}$ of blue light in Pro99 medium using 800 µM $NO_3^-$ as the sole N source for MIT0915 and MIT0917 and 100 µM $NO_2^-$ as the sole N source for MIT1214. Cell concentrations were then determined using flow cytometry on a Millipore Guava flow cytometer. Co-cultures were established by inoculating fresh medium with 2 x $10^6$ cells $mL^{-1}$ of each strain for a total initial cell concentration of 4 x $10^6$ cells $mL^{-1}$. Pure cultures were inoculated into fresh medium at 4 x $10^6$ cells $mL^{-1}$. Cultures were monitored daily by removing 0.5 mL of culture for flow cytometry and $NO_2^-$ concentration determination as detailed above. Daily samples for quantitative PCR were preserved by filtering 1 mL of culture onto a 25 mm 0.2 µm pore size polycarbonate filter under low vacuum, chasing with 2 mL of qPCR preservation solution (10 mM Tris pH=8, 100 mM EDTA, and 500 mM NaCl), and then transferring the filter to a 2 mL beadbeater tube prior to storage at -80ºC. After the initial culture had grown for 7 days, a subsample was transferred into fresh medium at a final cell concentration of 8 x $10^7$ cells $mL^{-1}$ (total cells). The initial transfer was monitored for 14 days and the second transfer was monitored for 8 days (i.e., until the cultures began to enter stationary phase as indicated by the daily change in cell concentrations).

**Quantitative PCR methods.** Primers targeting the *wcaK* gene of *Prochlorococcus* MIT1214 were designed using the NCBI Primer-BLAST tool. The primer set includes the forward primer, MIT1214_wcaK_283F (5'-GACTACTGCATTTTCGCTGGG-3') and the reverse primer, MIT1214_wcaK_402R (5'- ACCTTCAAAACCTCCAACACC).

Samples used to generate standard curves were acquired by growing MIT0915, MIT0917, and MIT1214 to late-exponential phase (approximately 8 x $10^7$ cells $mL^{-1}$), filtering 5 mL of culture onto a 25 mm 0.2 µm pore size polycarbonate filter under low vacuum, chasing with 3 mL of qPCR preservation solution (10 mM Tris pH=8, 100 mM EDTA, and 500 mM NaCl), and then transferring the filter to a 2 mL beadbeater tube prior to storage at -80ºC. Cell concentrations for each culture, at the time of sample filtration, was obtained through flow cytometry. Templates for both experimental cultures and standards were generated by thawing the filters on ice for 2 min, adding 650 µl of 10 mM Tris pH=8, and then beadbeating at 4800 rpm for 2 minutes. Following beadbeating to remove cells from the filter, 500 µl of the buffer was transferred to a 1.5 mL centrifuge tube and heated at 95ºC for 15 min to lyse cells. Templates for standard curves were generated by first diluting the resulting template solution to 5.4 x $10^5$ cells $\mu l^{-1}$ and then performing a serial dilution. All templates were stored at -80°C until use.

The MIT1214 *wcaK* assay was performed in 25 µl reaction volumes with 2.5 µl template and the following final concentrations of reaction components: 12.5 µl QuantiTect SYBR Green PCR Mix (Qiagen, Germantown, Maryland) and 0.5 µmol $L^{-1}$ of each forward and reverse primer. Using a CFX96 Thermocycler (Bio-Rad, Hercules, CA, USA), reactions were pre-incubated at 95°C for 15 min to activate the polymerase and then cycled (40 cycles) at 95°C for 15 s, 57°C for 30 s, and 72°C for 30 s. The MIT0915 and MIT0917 *narB* assays were performed similarly, except for annealing at 60°C for 30 s [2]. Amplification efficiencies were 85% for the MIT1214 *wcaK* assay, 90% for the MIT0915 *narB* assay, and 79% for the MIT0917 *narB* assay. Negative controls included MIT0915 and MIT0917 templates for the MIT1214 *wcaK* assay as well as MIT1214 templates for the *narB* assay; no amplification was observed in these negative controls.

**Metagenomic derived frequencies of LLI functional types.** Paired-end sequencing reads for samples obtained from the subsurface chlorophyll maximum layer at HOT and BATS [3] were annotated using kaiju 1.7.2 [4] and the MARMICRODB reference database of marine microorganisms [5, 6]:

```
kaiju -z 20 \
       -t $DATABASE/nodes.dmp \
       -f $DATABASE/MARMICRODB.fmi \
       -i $READDATADIR/$LIBRARY_1_trimmed.fq \
       -j $READDATADIR/$LIBRARY_2_trimmed.fq \
       -o $LIBRARY_marmicrodb_pairs_kaiju.out -v
```

Reads matching the taxonomic identifier for the LLI clade of *Prochlorococcus* were extracted from the paired-end sequencing reads using seqtk 1.3 (https://github.com/lh3/seqtk). Frequencies of LLI *Prochlorococcus* N assimilation genotypes were determined by further annotation of the taxonomically binned reads using kaiju and the CyCOG v6 database [7]. Reads that mapped to the *gyrB, narB*, *focA*, type I *nirA*, and type II *nirA* genes were enumerated and normalized to gene length. We assume that each of these genes are found in single copies in *Prochlorococcus* genomes based on their prevalence in the CyCOG v6 database [7]. Fractions of LLI *Prochlorococcus* that belonged to each of the 3 functional types (Fig. 4 in the main text) were resolved using the gene length normalized counts of N assimilation marker genes in each metagenome. Specifically, the abundance of MIT1214-like genomes was operationally defined as length-normalized counts of type I *nirA* genes less the length-normalized counts of *narB* genes. MIT0917-like genomes were defined as the length-normalized counts of type II *nirA* genes. MIT0915-like genomes were defined as the length-normalized counts of *narB* genes less the length-normalized counts of type II *nirA* genes. Each of these values was divided by the length-normalized counts of the *gyrB* gene, a single copy core gene in *Prochlorococcus*, in order to obtain the fraction of the LLI *Prochlorococcus* population represented by each functional type. Seasonality of the frequency of each functional type in LLI *Prochlorococcus* populations was assessed by binning these data into 4 subsets based on sample collection month – winter (January through March), spring (April through June), summer (July through September), and autumn (October through December).

**Table S1.** *Prochlorococcus* and *Synechococcus* strains used in this study.

| Strain | Genus | Subcluster/Clade | Inorganic N Assimilation | Original Reference | Rendered Axenic |
|---|---|---|---|---|---|
| MIT0915 | *Prochlorococcus* | Clade LLI | $NH_4^+$, $NO_2^-$, $NO_3^-$ | [2] | This Study |
| MIT0917 | *Prochlorococcus* | Clade LLI | $NH_4^+$, $NO_2^-$, $NO_3^-$ | [2] | This Study |
| MIT1214 | *Prochlorococcus* | Clade LLI | $NH_4^+$, $NO_2^-$ | This Study | This Study |
| SB | *Prochlorococcus* | Clade HLII | $NH_4^+$, $NO_2^-$, $NO_3^-$ | [8] | [1] |
| WH7803 | *Synechococcus* | Subcluster 5.1B; Clade V | $NH_4^+$, $NO_2^-$, $NO_3^-$ | [9] | [9] |
| WH8102 | *Synechococcus* | Subcluster 5.1A; Clade III | $NH_4^+$, $NO_2^-$, $NO_3^-$ | [9] | [9] |

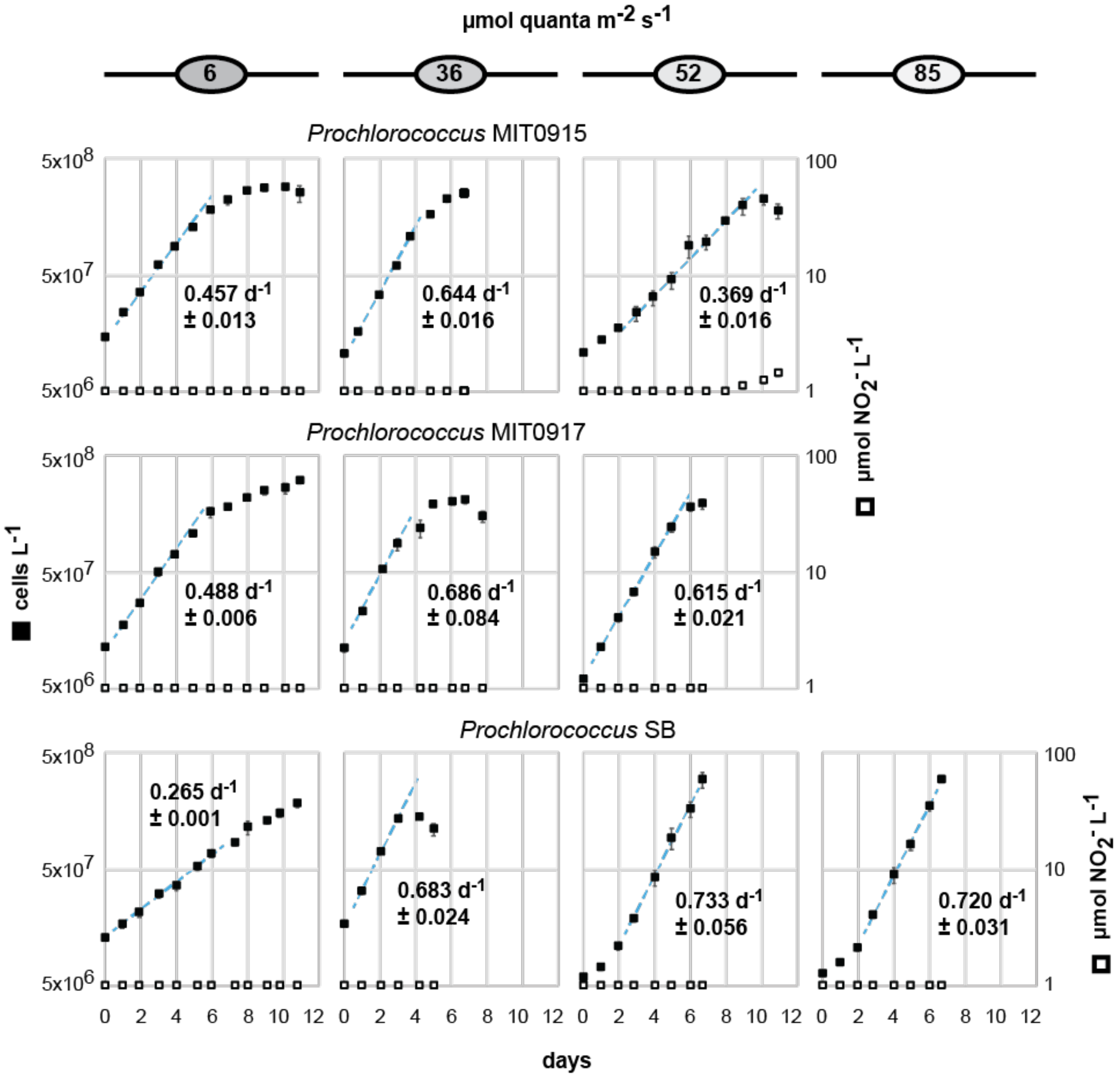


**Fig. S1.** Growth rates and extracellular $NO_2^-$ concentrations for batch cultures of *Prochlorococcus* grown on $NH_4^+$ as the sole N source over a range of light intensities. $NO_2^-$ concentrations below the dynamic range of the assay (< 1 μM $NO_2^-$) are plotted on the x-axis.

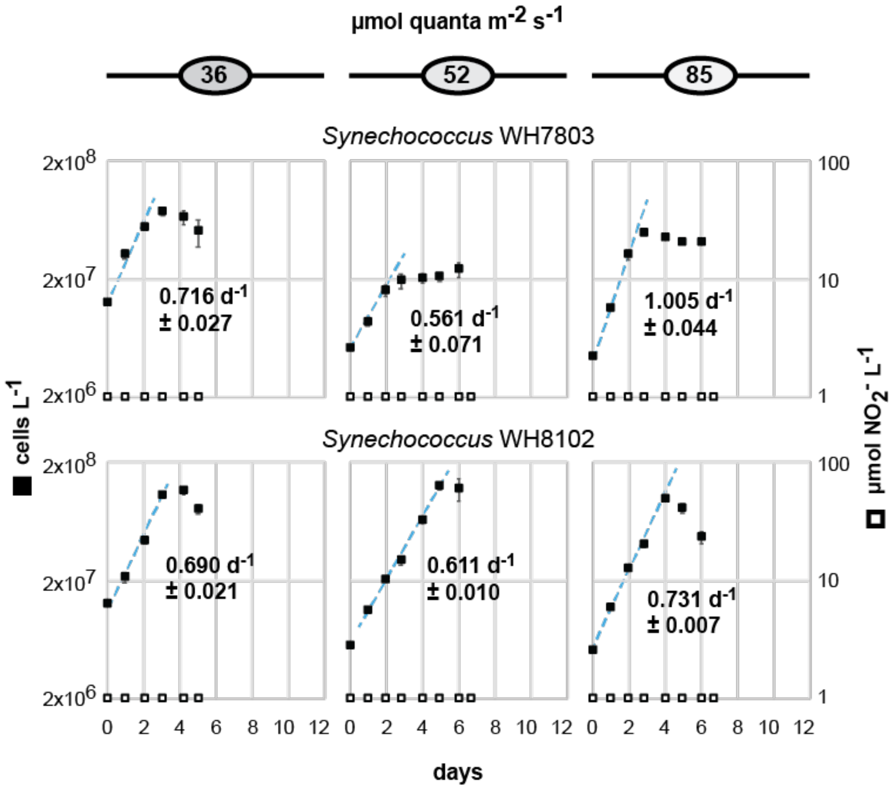


**Fig. S2.** Growth rates and extracellular $NO_2^-$ concentrations for batch cultures of *Synechococcus* grown on $NH_4^+$ as the sole N source over a range of light intensities. $NO_2^-$ concentrations below the dynamic range of the assay (< 1 μM $NO_2^-$) are plotted on the x-axis.
